# Arm position estimates derived from motor biases

**DOI:** 10.1101/2025.07.26.664535

**Authors:** Kevin Abishek, Hamza Muzammall, Hyunglae Lee, Christopher A. Buneo

## Abstract

Proprioception, including position sense, is critically important for normal sensorimotor and perceptual functioning but remains poorly understood. In the laboratory and clinic, arm position sense is typically assessed using perceptual tasks, e.g., via arm position matching or reporting the arm’s current position against a previously sensed one. Although such assessments provide important information about position sensing, they are incomplete in that they do not directly address the role of position sense in motor planning and control. Here, we used a combination of human psychophysical experiments, forward dynamic simulations, and mathematical optimization to reverse engineer arm positions (‘REAP’) used during motor planning. Subjects performed arm movements in a virtual environment under conditions where visual arm position cues were either aligned with corresponding somatosensory cues or were shifted prior to reach onset in one of four directions. Under shifted conditions, subjects exhibited characteristic ‘motor biases’, i.e., systematic deviations from ideal trajectories to the visually-cued targets. The arm positions used to plan movements under shifted conditions were obtained by minimizing differences between the experimentally induced motor biases and simulated motor biases. These REAP estimates largely conformed to predictions derived from previous perceptual experiments in that they were strongly influenced by the visual cues, and in a manner that was axially dependent. The results suggest that assessments of position sense derived from motor biases can be used to augment perceptual assessments or be used in lieu of them when perceptual reporting isn’t possible. In addition, the observed similarities between position estimates and weights derived from motor and perceptual tasks suggest that the brain’s perceptual and action systems use similar mechanisms to deduce arm position from somatosensory and visual cues.

## 1 Introduction

Proprioception, often referred to as the “sixth sense”, comprises the senses of body position (aka ‘position sense’), movement (aka ‘kinesthesis’), and effort/force/heaviness (Proske & Gandevia, 2012). These senses contribute not only to the formation and maintenance of the body image, which contributes to the sense of limb ownership, but also to the body schema, which is important for motor control. Proprioceptive loss or impairment, i.e., proprioceptive dysfunction, can result from several conditions affecting the nervous system, including traumatic brain injury, stroke, Parkinson’s disease, Huntington’s disease, autism spectrum disorder, and diabetes. Proprioceptive dysfunction can also result from injuries and surgeries affecting the joints. Proprioceptive dysfunction can have dramatic effects on the body image, as in the case of somatoparaphrenia, where one denies ownership of a limb or an entire side of one’s body (Feinberg & Venneri, 2014), but also on the body schema, impairing the planning and control of limb and body movements (Ghez et al., 1995; Gordon et al., 1995). Both manifestations of proprioceptive dysfunction can adversely affect the performance of activities of daily living and negatively impact quality of life.

Position sensing remains enigmatic, despite the fact that it is important for both normal perceptual and sensorimotor function. One major factor contributing to this general lack of understanding is that methods for characterizing position sense primarily address its perceptual components. In the case of the arm, these methods include, among others, position matching (i.e., replicating the position of one arm with the other) reporting the arm’s current position against a previously visited one, pointing to the position of the unseen (tested) arm, and reporting the arm’s position with respect to an extrinsic landmark (Dukelow et al., 2010; Fuentes & Bastian, 2010; Klein et al., 2018; Rincon-Gonzalez et al., 2011; Simo et al., 2014; Wilson et al., 2010). Although such assessments can provide important information about arm position sense, they do not directly address position estimation in the context of motor planning and control and therefore likely provide only an incomplete picture of this critical function. In fact, observations that post-stroke recovery of motor function is not strongly correlated with recovery of proprioceptive abilities (Dukelow et al., 2012) may be at least partially due to the fact that existing assessment methods are largely perceptual and not motor in nature. A complete characterization of arm proprioception, therefore, should not only include methods that more effectively probe proprioception’s role in conscious awareness of position and movement, but also its role in motor planning and control, the details of which operate at a largely unconscious level.

Arm motor planning can be envisioned as a series of computations that gradually convert desired movement goals into appropriate spatiotemporal patterns of muscle activations. Arm position information is required at two distinct stages in this process. For example, when reaching for a cup (Fig. 1), arm position is initially used to determine the required movement vector, i.e., the direction and distance the hand must move to reach its goal location. In situations where the hand can be seen as well as felt, this vector planning (VP) stage is thought to depend predominantly on vision. At a later stage, the movement vector is subsequently transformed into a hand trajectory, i.e., a time-varying sequence of hand positions, which is later transformed via inverse modeling (IM) into a time-varying sequence of joint-based motor commands. In contrast to the VP stage, the IM stage requires information about the arm’s configuration, i.e., the angles of the shoulder and elbow joints, which is provided predominantly by muscle, joint and cutaneous afferents of the somatosensory system (Sober & Sabes, 2005). Importantly, errors in sensing the position of the arm during the VP stage, IM stage, or both would be expected to adversely affect the motor planning process.

**Figure 1.**
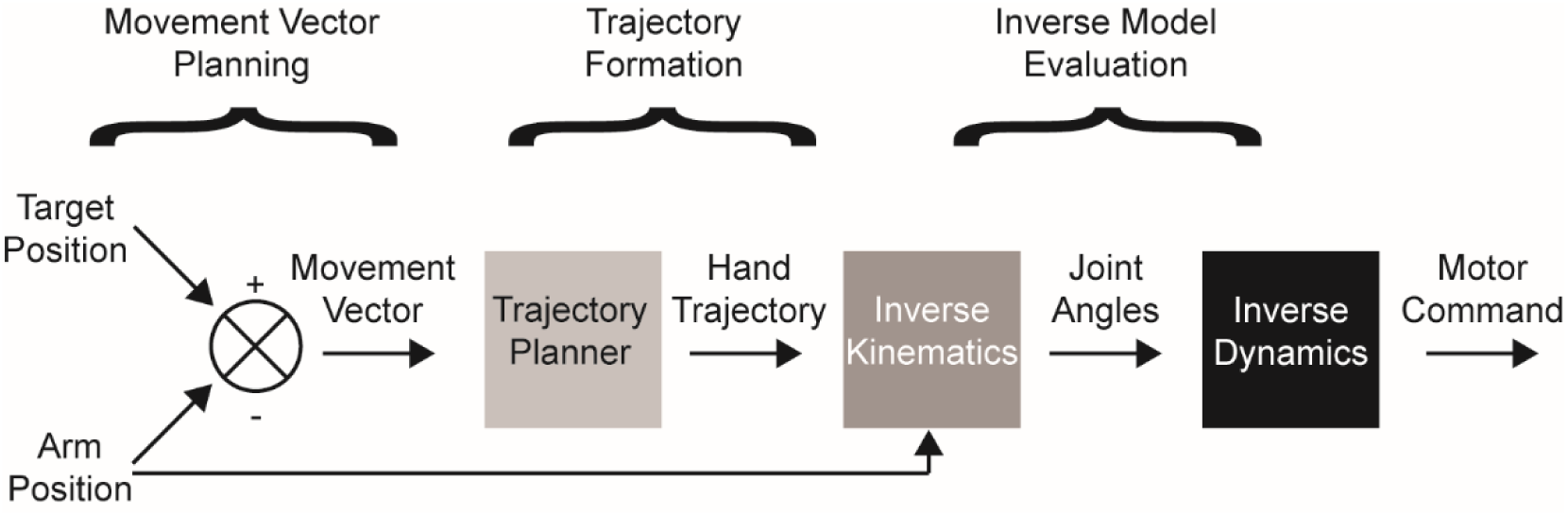
Schematic diagram of computations involved in planning reaching movements.

Sensing the position of the arm during motor planning can be conceived of as a Bayesian estimation process, where the brain attempts to determine the arm’s veridical position by combining a ‘sensory estimate’ derived from somatosensory and visual proprioceptive signals with prior knowledge and a ‘motor estimate’ derived from efference copy and a forward model of the arm (Körding & Wolpert, 2006). We will refer to the outcome of this process simply as the ‘position estimate*’*. Sensor noise and other factors can interfere with this process, leading to position *misestimation*, which can be characterized by its bias (the average direction and distance of estimates with respect to the arm’s veridical position; cross in Fig. 2A) and its variance (the similarity of multiple estimates to each other; dots and ellipse in Fig. 2A). Small, experimentally induced discrepancies between the seen and felt positions of the arm (<7 cm), though not consciously perceived (Hsiao et al., 2022), nonetheless result in biased position estimates, due to the differential weighting of visual and somatosensory cues during multisensory integration. Moreover, previous studies have shown that arm position biases are associated with distinct patterns of motor bias that manifest as systematic deviations in initial movement directions (Rossetti et al., 1995; Sober & Sabes, 2003, 2005; Vindras et al., 1998). This suggests that quantification of motor biases could, in principle, be used to infer position estimates that were used during motor planning.

**Figure 2.**
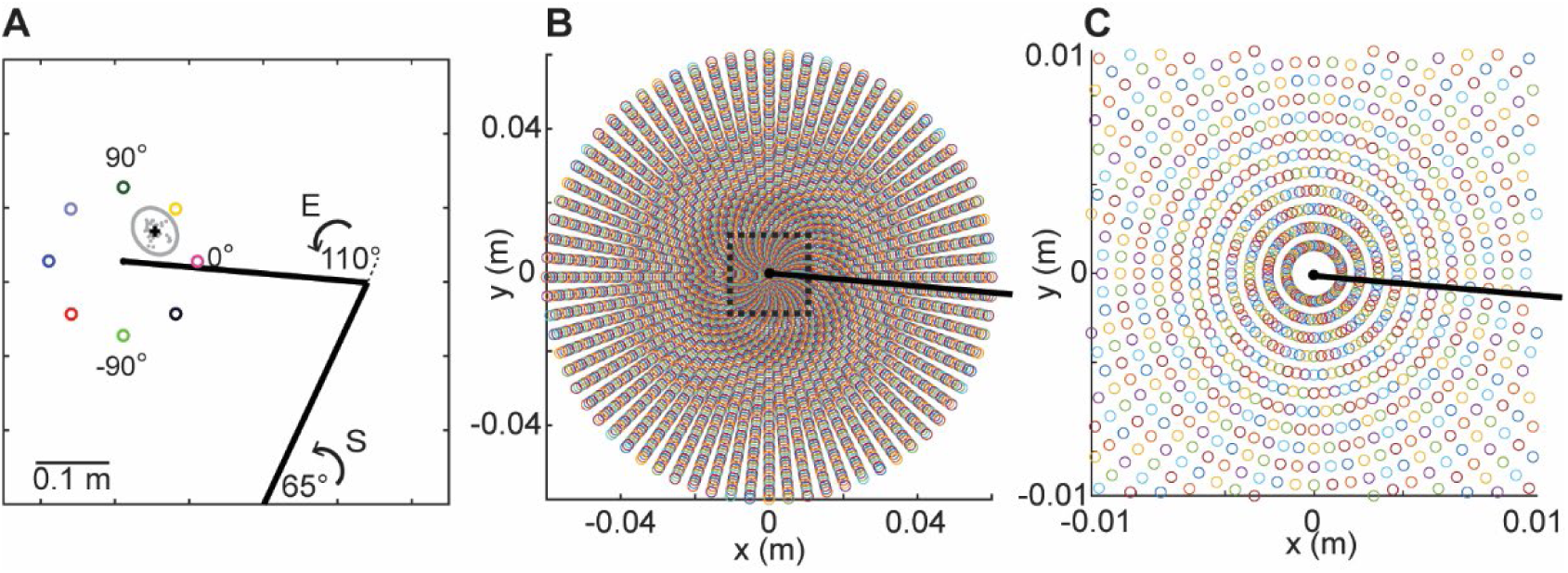
Simulated arm position biases. A. Initial arm position and target positions used in the simulations. Grey ellipse and points: distribution of biased arm positions with a mean centered at 45° and 0.04 m from the endpoint of the arm. B. Distribution of position biases (N=5,184) used to generate motor biases. Black line: forearm. C. Enlarged depiction of the biased arm positions indicated by the dotted square in B.

Here we describe a method to reverse engineer arm position (‘REAP’) estimates and sensory weights used during motor planning, enabled in part by analyses of simulated and experimentally induced motor biases. Based on previous studies (Sober & Sabes, 2003; van Beers et al., 1998, 1999; van Beers et al., 2002) we hypothesized that position estimates would generally be strongly biased by vision, but that the extent of this bias would depend upon the axis along which visual and somatosensory cue integration occurs. That is, estimates would be more strongly biased by vision when visual arm position cues were shifted laterally relative to corresponding somatosensory cues, than when these shifts occurred along an anteroposterior axis. We first describe the results of forward dynamic simulations designed to expand upon existing knowledge regarding the relation between arm position bias and motor bias and to aid in interpreting experimentally induced movement biases. More specifically, we show how position biases in different directions and magnitudes are expected to differentially affect the VP and IM stages of motor planning, and how these position biases lead to distinct biases in not only planned movement directions but planned movement distances and speeds as well, which has not previously been demonstrated. We then describe the results of behavioral experiments designed to quantify the relation between position biases and motor biases in healthy adult subjects. Lastly, we show how experimental motor biases and simulated motor biases can be used to reverse engineer arm positions that were used during motor planning.

## 2 Materials and Methods

### 2.1 Simulations

Previous studies have used forward dynamics modeling and simulations to gain insights into the effects of position bias and variance on motor planning and control, and a similar approach was used here (Buneo et al., 1995; Shi & Buneo, 2009; Sober & Sabes, 2003, 2005). Two joint (shoulder and elbow) arm movements in a horizontal plane were simulated using custom MATLAB code (The MathWorks Inc., Natick, MA). Representative limb segment lengths, limb masses, moments of inertia and centers of mass were obtained from previous simulation studies (Scheidt et al., 2005). A fixed initial arm posture of 65° shoulder horizontal adduction and 110° elbow flexion was used for all simulations (Fig. 2A). Assuming upper arm and forearm lengths of 0.33 m, these joint angles corresponded to an arm endpoint position that was ∼0.3 m anterior and 0.2 m medial to the shoulder joint, similar to the initial position of the hand in the experiments (see below). Simulated ‘baseline’ movements were made to targets at a distance of 0.1 m from the starting position and were 0.9 s in duration, which is consistent with the target distances and average movement durations used in the experiments. Simulations involved a combination of inverse and forward modeling and were entirely feedforward; visual and somatosensory feedback control were not simulated. Inverse modeling was used to determine the endpoint trajectories, joint angles, and joint torques consistent with movements in various directions and distances. We first produced idealized trajectories in Cartesian endpoint coordinates under the assumptions that hand paths are rectilinear and tangential speed profiles are bell-shaped (Hogan & Flash, 1987; Morasso, 1981; Soechting & Lacquaniti, 1981). Hand positions along the planned trajectories were then converted to time-varying angular positions at the shoulder and elbow using standard trigonometric equations. After numerical differentiation of the angular positions, shoulder and elbow torques were calculated using inverse dynamics (Hollerbach & Flash, 1982).

After determining the endpoint trajectories, joint angles, and joint torques associated with different planned directions and distances, movements were generated by inverting the procedure described above. Joint torques were used to derive movements in joint coordinates by simultaneously solving for the shoulder and elbow angular positions and velocities using a fourth-order Runge-Kutta method with a variable time step. Trajectories in endpoint coordinates were obtained by transforming the joint angular motions into hand paths.

These baseline simulation procedures were modified to characterize the motor biases that are expected to result from different arm position biases. Note that although these position biases were assumed to result from the integration of sensory and motor cues, we did not explicitly simulate this integration. To simulate position biases, on each run of the simulations, we introduced a discrepancy between the arm position used to calculate the required joint torques for a desired movement and the information used to solve for the joint angles and velocities at each time step. Arm position biases were introduced in 72 directions (every 5°) with respect to the initial arm endpoint position and at each of 72 magnitudes (from 0.0001m to 0.07 m at ∼0.0001 m intervals; Fig. 2B, C). For each position bias (N=5184), arm movements were simulated in eight directions separated by 45°. To be consistent with the experimental data analyses, the beginning and end of each movement were defined as the points at which the tangential hand speed first exceeded and then fell to 5% of its peak value.

Initial directional biases (DIRB) were quantified by calculating the angular differences between the position vectors specifying the endpoint of the arm at 40% of peak tangential speed and the vectors connecting the initial position endpoint to the targets (Fig. S1). This percentage of peak speed was chosen to coincide with that used to quantify directional biases in the experiments, which was consistent with previous behavioral studies (Sober & Sabes, 2003). For a given movement direction, initial distance biases (DISB) and speed biases (SB) were calculated by taking the difference in displacement and speed (at 40% of peak tangential speed) between biased and unbiased (i.e., baseline) arm position conditions.

As described in Results, introducing arm position biases affected both the VP and IM stages of the simulations, resulting in changes in planned movement directions and distances. Changes in planned movement distances also resulted in concomitant changes in movement speeds. More precisely, given that movement durations were fixed, peak movement speeds scaled linearly with planned movement distances, an assumption that appears reasonable for the relatively small movements employed here (Gordon, Ghilardi, Cooper, et al., 1994).

### 2.2 Subjects

Twenty (20) individuals (nine males, 11 females) without prior history of neurological, psychiatric, or orthopedic disorders affecting the right arm participated in these experiments. All individuals participated in Experiment 1, while six of these same individuals also participated in Experiment 2. Participants declared themselves to be right-handed, but their handedness was also assessed using the Edinburgh Handedness Inventory -Short Form (Veale, 2014). All subjects reported having normal or corrected-to-normal vision. Informed, written consent was obtained from all participants in accordance with the Declaration of Helsinki, and all experimental procedures were approved by the Arizona State University Institutional Review Board.

### 2.3 Apparatus and Behavioral Task

Experimental data were collected using a KINARM End-Point Lab (BKIN Technologies Ltd., Kinston, ON, CA). This apparatus consists of a planar robotic arm with a graspable handle, 1kHz position sampling, and a 2D virtual/augmented reality (VR/AR) display. All participants used their right (dominant) arm to perform the task. Subjects were instructed to follow the visual cues that were presented and to reach quickly and accurately to the cued targets, using a single uncorrected movement. No other specific instructions were provided, including regarding movement speed. Movements were initiated from a fixed starting position that was located on the body midline and approximately 0.3 m from the frontal plane of the body. Due to space limitations imposed by the KINARM system, the plane containing the upper arm and forearm differed in orientation from that used in the simulations, i.e., it was oriented more vertically rather than horizontally. Targets were located at a fixed distance of 0.1 m from the starting position and in eight directions spaced 45° apart, i.e., in a ‘center-out’ target arrangement. Vision of the real arm was blocked by the apparatus, but a virtual visual cue representing the endpoint of the arm/robot handle was intermittently provided in the form of a 0.025 m radius white filled circle, as described below.

#### Experiment 1

Subjects initially performed a series of practice trials involving center-out movements performed with the visual hand position cue aligned with the robot handle and provided throughout the trials, in order to enforce the idea that this cue represented their hand position. At the beginning of these trials, subjects grasped the robot handle, which was then moved by the robot controller to the starting position. After holding at the starting position for 0.5-1 second, a visual target cue was then presented in the form of a 0.025 radius filled red circle, which also served as the ‘go’ cue for the movement. Trials ended when tangential hand speed fell below 0.001 m/s. At the end of each trial, the robot controller then moved the handle back to the starting position for the next trial. Three trial were performed for each of the eight target directions, for a total of 24 trials. Additional blocks of practice trials were provided as needed by the subjects.

On subsequent experimental trials, the visual hand position cue was initially absent. Subjects grasped the robot handle, which was again moved passively to the starting position. After holding at the starting position for 0.5-1 second, the visual hand position cue was displayed for a variable period of 0.5 −1.5 seconds, either at the same position as the handle or shifted 0.04 m medially, laterally, anteriorly, or posteriorly with respect to the handle. Following this, the visual target position cue was presented, cueing the movement. Once the handle of the robot moved 0.005 m from its initial position, the visual hand position cue was extinguished, and the remainder of the movement was performed without hand visual feedback. At the end of a trial, the robot controller then moved the handle back to the starting position for the next trial.

Importantly, several steps were taken to minimize the likelihood of adaptation and recalibration/reweighting of sensory cues. First, as indicated above, visual arm position cues were only shown at the starting position and were removed after the robot handle had moved a very short distance from the starting position, i.e., very minimal online visual feedback was provided. As a result, subjects were also prevented from seeing their final hand positions with respect to the targets, i.e., no knowledge of results was provided. Lastly, visual shifts in different directions and magnitudes were pseudo-randomly interleaved on a trial-by-trial basis, thereby further minimizing the likelihood of motor adaptation.

#### Experiment 2

Six subjects participated in this experiment, which was designed to determine the extent to which hand positions derived from the REAP procedure in Experiment 1 reflected subjects’ actual estimated hand positions. In contrast to Experiment 1, only two visual hand position cue shifts were used: a zero-shift condition and a 0.04 m lateral shift condition. In addition, and unbeknownst to the subjects, all targets in this experiment were shifted laterally by an amount equal to the average REAP-derived hand position estimate in the lateral shift condition in Experiment 1 (i.e., 0.033 m lateral). The rationale for this was as follows: if the REAP-derived positions truly reflected subjects’ estimated hand positions in the lateral shift condition, then shifting the targets to an equal degree would result in subjects planning movements with minimal directional biases, as the required movement vectors would be largely identical to those required in the zero-shift condition. Other than the reduced number of conditions and the shifted positions of the targets, all other experimental procedures and parameters were identical to those in Experiment 1.

### 2.4 Data Analysis

Hand position data on each trial were smoothed using a double-pass, zero-lag filter with a cutoff frequency of 3dB. Hand velocity was calculated directly from the KINARM robot joint kinematics. As in the simulations, the points at which the tangential hand speed first exceeded and then fell to 5% of its peak value were used to identify the beginning and end of each movement. The hand position corresponding to 40% of peak speed was used to estimate motor biases. This represented a trade-off between minimizing the influence of online proprioceptive feedback and minimizing the effects of noise in the speed profiles (Sober & Sabes, 2003, 2005).

Importantly, motor biases were calculated using a slightly different procedure than that used for the simulations. In simulation, no directional biases are generated under ‘baseline’ conditions, i.e., in the absence of an imposed arm position bias. As a result, directional biases under conditions of imposed biases can therefore be computed with respect to either the baseline movement directions or the target directions, as these did not differ by any appreciable amount. However, human subjects have previously been shown to exhibit characteristic ‘baseline’ biases in initial movement directions even in the absence of an imposed position bias (Wang, 2024). These directional biases could result from biased estimates of target position, inherent biases in somatosensory and/or visual estimates of arm position, and/or other factors related to motor planning and execution. Therefore, to minimize the influence of these ‘other’ processes, each subject’s experimental biases in direction, distance, and speed were computed with respect to their own initial directions and speeds in the baseline (no visual shift) condition.

### 2.5 REAP Procedure

Our main focus was on using directional biases to infer subject position estimates. Although we could have used both initial distance and speed biases as well, these movement parameters tended to covary in the experiments (Fig. S3). This observation, combined with the knowledge that motor actions during planning appear to be represented in terms of their directions and speeds (Moran & Schwartz, 1999), led us to a secondary focus on speed biases only. Note that, due to the relatively large variation in peak movement speeds across subjects, speed data were z-score normalized for the optimization analyses.

For each subject, experimental motor biases (directional or speed) associated with a given visual shift condition were compared to each of the 5184 patterns of simulated motor biases to determine the ‘best-fitting’ position bias. More specifically, for directional biases, position estimates were determined by finding the direction and magnitude of the simulated arm position bias that minimized the following error function:

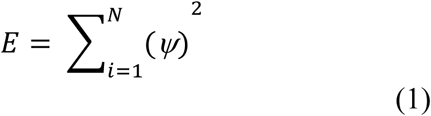

where *ψ* is the angle between experimental and simulated movement directions (at 40% of peak speed) for each of *N* = 8 target directions.

An analogous procedure was used for speed biases, with the exception that *ψ* represented the difference between experimental and simulated movement speeds at 40% peak speed. For both directional and speed biases, the minimization procedure was accomplished in two steps. First, we fit a surface to these differences in order to obtain an objective function to minimize. Both cubic interpolation and LOESS fitting procedures (implemented in MATLAB) were examined. After obtaining an objective function, an unconstrained minimization routine employing a quasi-Newton algorithm (‘fminunc’; MATLAB’s Optimization Toolbox) was executed. Results were obtained for both the cubic interpolation and LOESS fits, though the former produced results that were somewhat more consistent across subjects. As a result, only the cubic interpolation generated results are presented here. Corresponding visual weights were determined by projecting the vectors defining the estimated arm positions derived from the REAP procedure onto those defining the visual shifts and dividing the result by the shift magnitude. Somatosensory weights were then taken to be 1-WV.

### 2.6 Statistical Analyses

All statistical analyses were conducted in SPSS 25 using an alpha of 0.05. Position estimates and visual weights associated with each shift were tested for normality using the Shapiro-Wilk test. T-tests were conducted on the x and y coordinates of the position estimates to determine if they differed significantly from zero. Differences in visual weights across the different shifts were then assessed using a one-factor repeated measures analysis of variance (ANOVA), followed by multiple comparisons using the Bonferroni procedure.

## 3 Results

### 3.4 Simulations

In simulation, VP errors led to motor biases that varied strongly with target direction. Figure 3A shows a schematic representation of a simulation where an arm movement was planned to a single, anteriorly located target using a biased arm position estimate (direction: 0°, magnitude: 0.04 m). In general, position biases of a given direction and magnitude resulted in an equal and opposite shift in planned movement endpoints. Thus, in the example in Fig. 3A, where arm position was biased anteriorly and to the right of the arm’s veridical position, the planned movement endpoint shifted posteriorly and to the left. This resulted in an incorrectly planned movement vector, i.e., one directed anteriorly and to the left of the hand’s veridical position. Figures 3B-D show the consequences of this position bias more generally, i.e., for ‘center-out’ movements planned to eight targets located on a circle around the veridical hand position. For all targets, planned movement endpoints shifted opposite the shift in the biased arm position, i.e., posteriorly and medially (Fig. 3B). These shifts in planned endpoints had differential effects on planned movement vectors (Fig. 3C), resulting in target-dependent directional biases (DIRB), as well as target-dependent distance biases (DISB). Note that the DIRB and DISB as shown here are for illustrative purposes only; for the REAP procedure these biases were calculated using the position corresponding to 40% of peak speed from the fully simulated trajectories (not the planned vectors). Targets located along an axis collinear with the direction of position bias (i.e., 45° and 135°) were associated with relatively large DISB but small DIRB, while orthogonal targets demonstrated large DIRB and relatively small DISB. Targets located along other axes showed intermediate effects. Both DIRB and DISB varied sinusoidally with target direction (Fig, 3D; see also (Sober & Sabes, 2003)), with the two bias patterns exhibiting a phase shift of approximately 90°.

**Figure 3.**
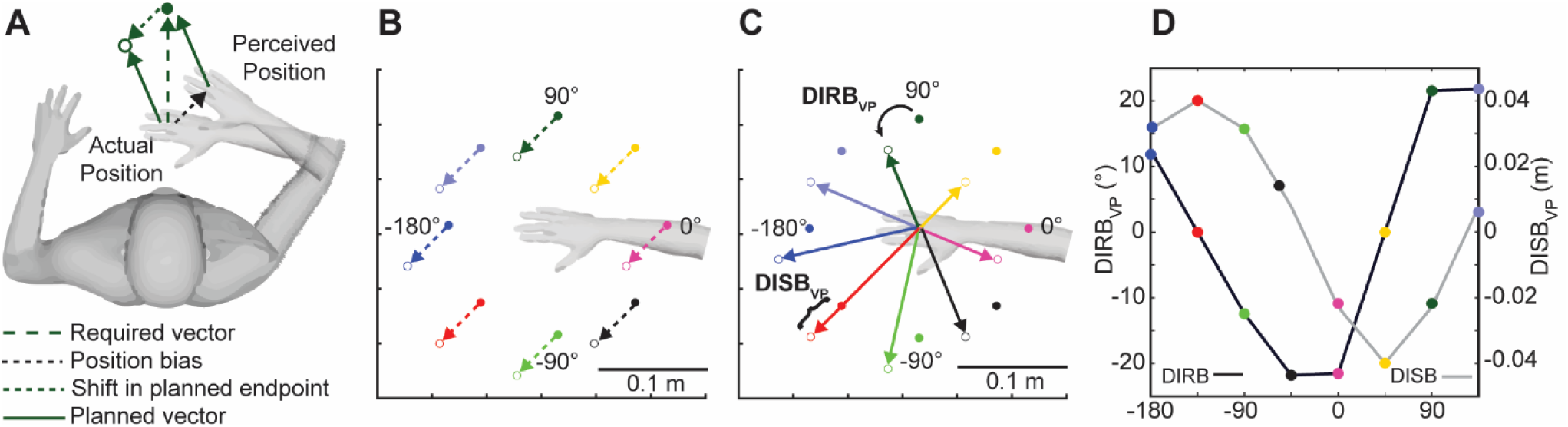
Motor biases resulting from vector planning errors. A. Illustration of the effects of an arm position bias of 45°, 0.04 m, on the planned movement vector and movement endpoint for a single target. B. For the same arm position bias, shifts in planned endpoints for eight targets centered on the unbiased endpoint of the arm. C. Concomitant changes in planned movement vectors to each target. DIRB_VP_: Directional bias arising from vector planning error. DISB_VP_: Distance bias arising from vector planning error. D. DIRB_VP_ and DISB_VP_ as a function of target direction. Vector planning errors result in motor biases that are highly direction-dependent.

In contrast to VP errors, IM errors resulting from position biases led to motor biases that were much less dependent on target direction. Figure 4 shows simulation results using the same position bias as in Fig. 3 but with no VP errors included. In other words, movement directions and distances were specified correctly, but were otherwise planned and executed using a biased arm configuration. First, Fig. 4A shows the full simulated trajectories when arm position was unbiased. Under these ‘baseline’ conditions, no VP or IM errors were generated, and as a result, simulated movements exhibited little to no motor biases. In contrast, when arm position was biased (Fig. 4B), the resulting movements were also biased, but in a different manner than when VP errors are present. Without the inclusion of VP errors, DIRB were greatly reduced and were more consistent in direction across target directions, in this case rotating consistently counterclockwise. In addition, DISB were also much smaller and largely independent of target direction. These trends can also be observed in Fig. 4C, which shows that DIRB were approximately 10°, regardless of target direction, while DISB were even smaller but also relatively consistent in magnitude across targets.

**Figure 4.**
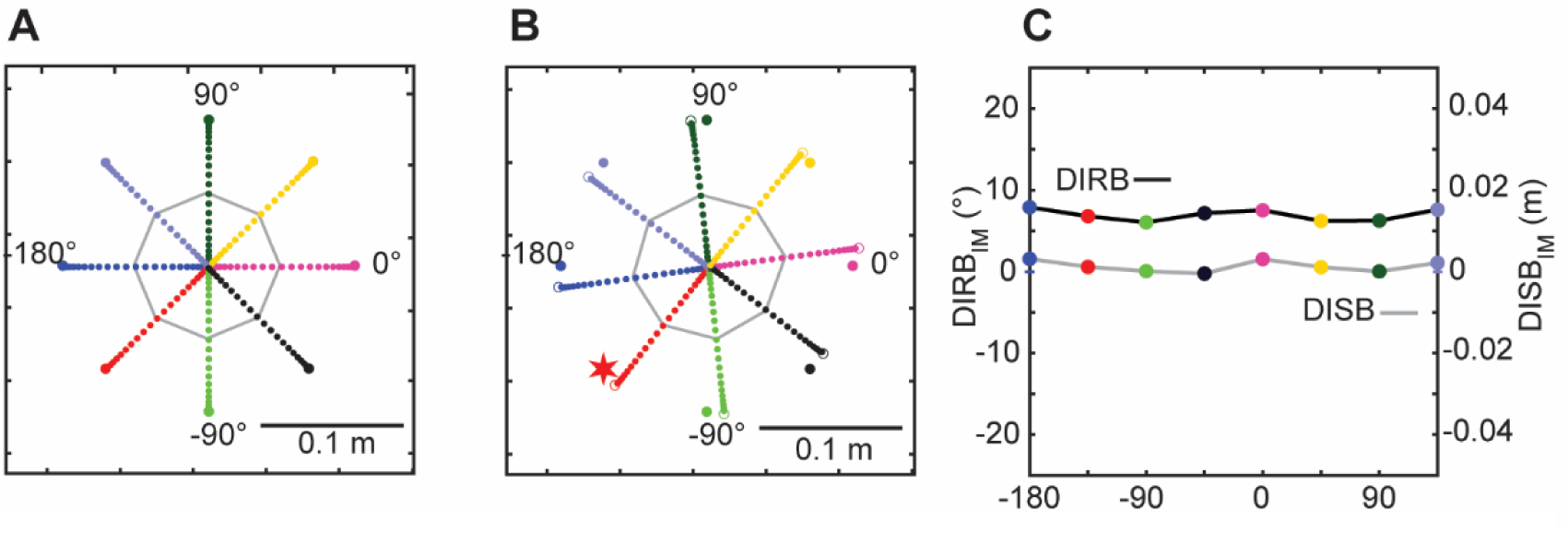
Motor biases resulting from inverse modeling errors. A. Movement trajectories when arm position is correctly estimated, i.e., in the absence of vector planning and inverse modeling errors. B. Movement trajectories in the presence of an arm position bias of 45°, 0.04 m. C. DIRB_IM_ and DISB_IM_ as a function of target direction. Inverse modeling errors resulted in biases that were relatively consistent in sign and magnitude across target directions.

Simulated directional biases varied with the direction (and magnitude) of arm position bias, but in a manner that also depended on initial arm configuration. Figure 5 shows patterns of DIRB for two different arm configurations (rows) and four different directions of arm position bias (colored lines). DIRB arising from VP errors alone, IM errors alone, and the combination of both factors are shown in columns B through D, respectively. As also illustrated in Fig. 3, Fig. 5B shows that DIRB arising from VP errors alone varied strongly with target direction. However, the specific pattern of DIRB across target directions depended strongly on the direction of arm position bias, with biases separated by 90° resulting in patterns of movement direction bias that were also phase shifted by ∼90°.

**Figure 5.**
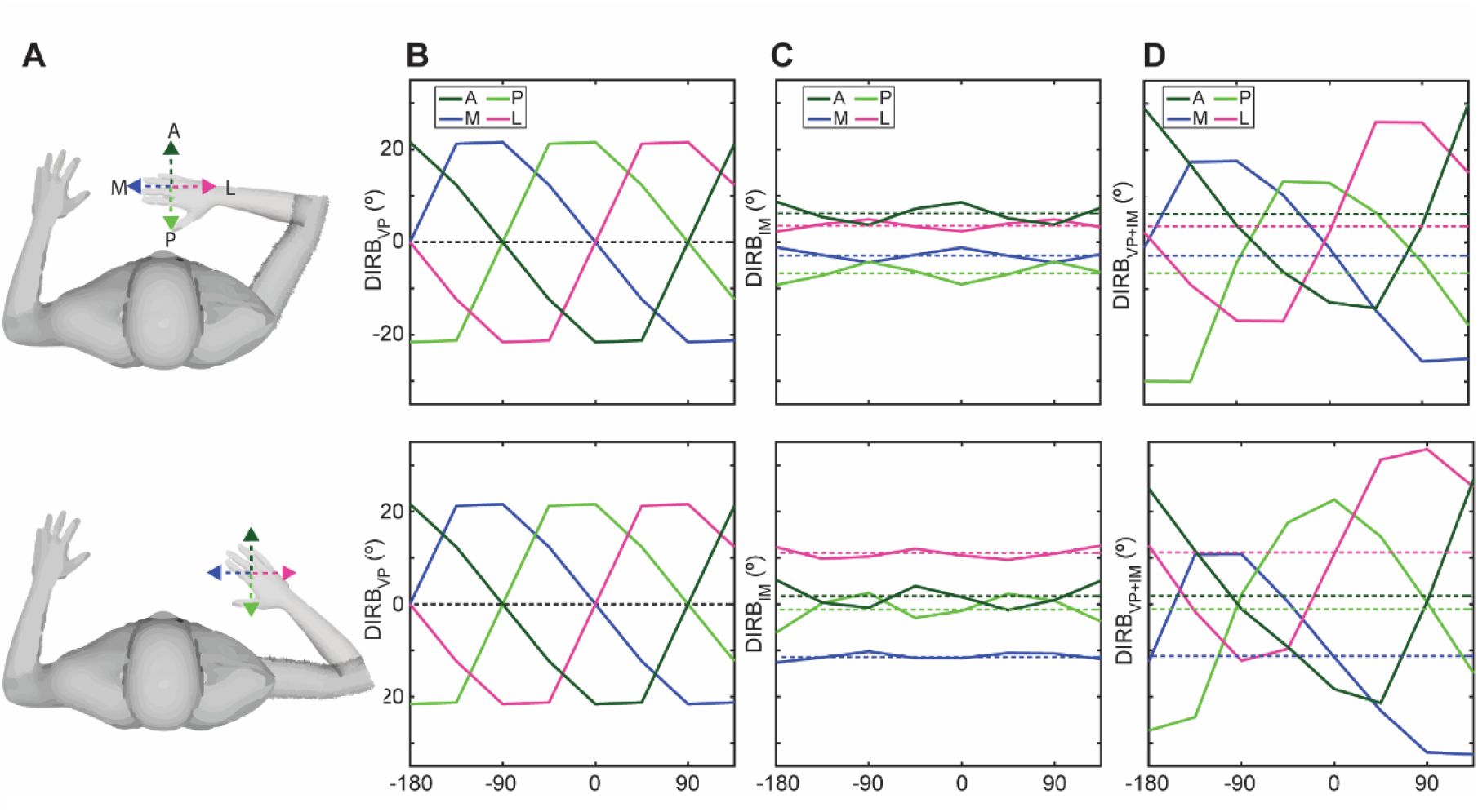
Effects of arm posture and direction of arm position bias on initial movement directions. A. Simulated arm postures and position biases. A bias magnitude of 0.04m was used for each direction. B. DIRB resulting from VP errors for the two arm postures. C. DIRB resulting from IM errors. D. Combined effect of VP and IM errors. DIRB_IM vary_ with arm posture but DIRB_VP_ do not. The combined effect of IM and VP errors results in a DC offset of the DIRB_Vp+IM_ vs target direction curves.

Regarding effects of arm configuration, a comparison of the plots in the different rows reveals that, if target directions remain fixed with respect to the initial hand position, VP-induced DIRB do not vary with configuration. In contrast, Fig. 5C shows that IM induced DIRB varied only weakly with target direction but exhibited different levels of DC offset depending on arm position bias. Moreover, in contrast to those induced by VP errors, IM induced DIRB varied with arm configuration, with the magnitude of DC offset either increasing or decreasing depending on the direction of arm position bias. As a result, when VP and IM errors were combined, DIRB varied to some degree with arm configuration (Fig. 5D), largely due to the arm configuration dependence of IM-induced DIRB.

Simulated arm position biases not only led to biases in movement direction and movement distance, but also to biases in movement speed. Figures 6A-B show for the same example arm position bias illustrated in Figs. 3 and 4 (direction: 0°, magnitude: 0.04 m), the movement paths, DIRB, and DISB that resulted from combined VP and IM errors. As previously shown for VP errors alone, DIRB and DISB varied strongly with target direction under these conditions. Regarding movement speeds, we assumed that peak movement speed scaled linearly with movement distance. This was a reasonable assumption based on previous studies (Gordon, Ghilardi, Cooper, et al., 1994) but was also largely borne out in our own experimental data (Fig. S3). A consequence of this assumption was that when planned movement directions and distances were altered in the presence of arm position bias (Fig. 6A, B), peak movement speeds changed as well. Figure 6C shows, for the −135° target direction, how movement speed varied with movement distance when arm position was biased compared to when it was not. Similar to Fig. 2, the SB shown here is only for illustrative purposes; for the REAP procedure, speed biases were calculated using the position corresponding to 40% of peak speed, rather than 100% as shown here. Under biased arm position conditions, peak speed occurred later and was greater in magnitude, leading to a ‘bias’ in peak speed. Figure 6D shows that, similar to DIRB and DISB, speed bias (SB) was strongly target direction dependent, matching very closely the pattern observed for DISB.

**Figure 6.**
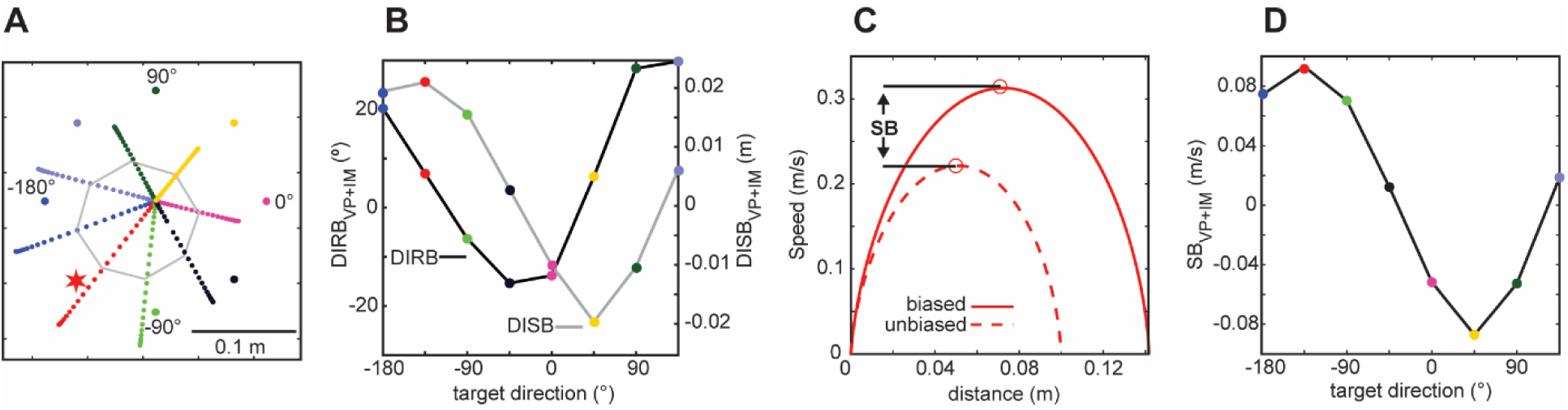
Speed biases. A. Movement trajectories generated with a 45°, 0.04m arm position bias, with resulting planning errors. B. DIRB and DISB as a function of target direction. C. Movement speed vs movement distance for the −135° target direction. Also shown is the expected speed-distance curve under the unbiased condition. Speed bias (VB) is defined as the difference in speed between unbiased and biased conditions. For illustrative purposes, peak speed was used in this example. VB as a function of target direction.

### 3.5 Experimentally induced directional biases

As predicted by the simulations, experimental DIRB varied with target direction, with the pattern of variation being highly dependent upon visual shift direction. Figure 7 shows, for a single representative subject, hand paths and DIRB associated with the ‘no visual shift’ condition (A), as well as those associated with visual shifts in four directions (B-E). As previously demonstrated by several groups (Ghilardi et al., 1995; Gordon, Ghilardi, Cooper, et al., 1994; Vindras et al., 1998), even in the no shift condition (A), i.e., in the absence of an imposed arm position bias, patterns of target dependent directional bias can still be observed, peaking in this example at target directions of −90° and +90°. Regarding the visual shift conditions (B-E), as previously shown for medial and lateral shifts (Sober & Sabes, 2003, 2005), DIRB varied in an approximately sinusoidal fashion with target direction and in a manner that was highly dependent upon shift direction. Figure 8 shows, for the same subject, movement speeds and distances (A) and their biases (B-E) under the same visual shift conditions. Speed and distance biases were generally more variable than DIRB but also varied in an approximately sinusoidal manner with target direction. Notably, these patterns of variation were phase-shifted relative to their associated DIRB, as predicted by the simulation results (Fig. 6).

**Figure 7.**
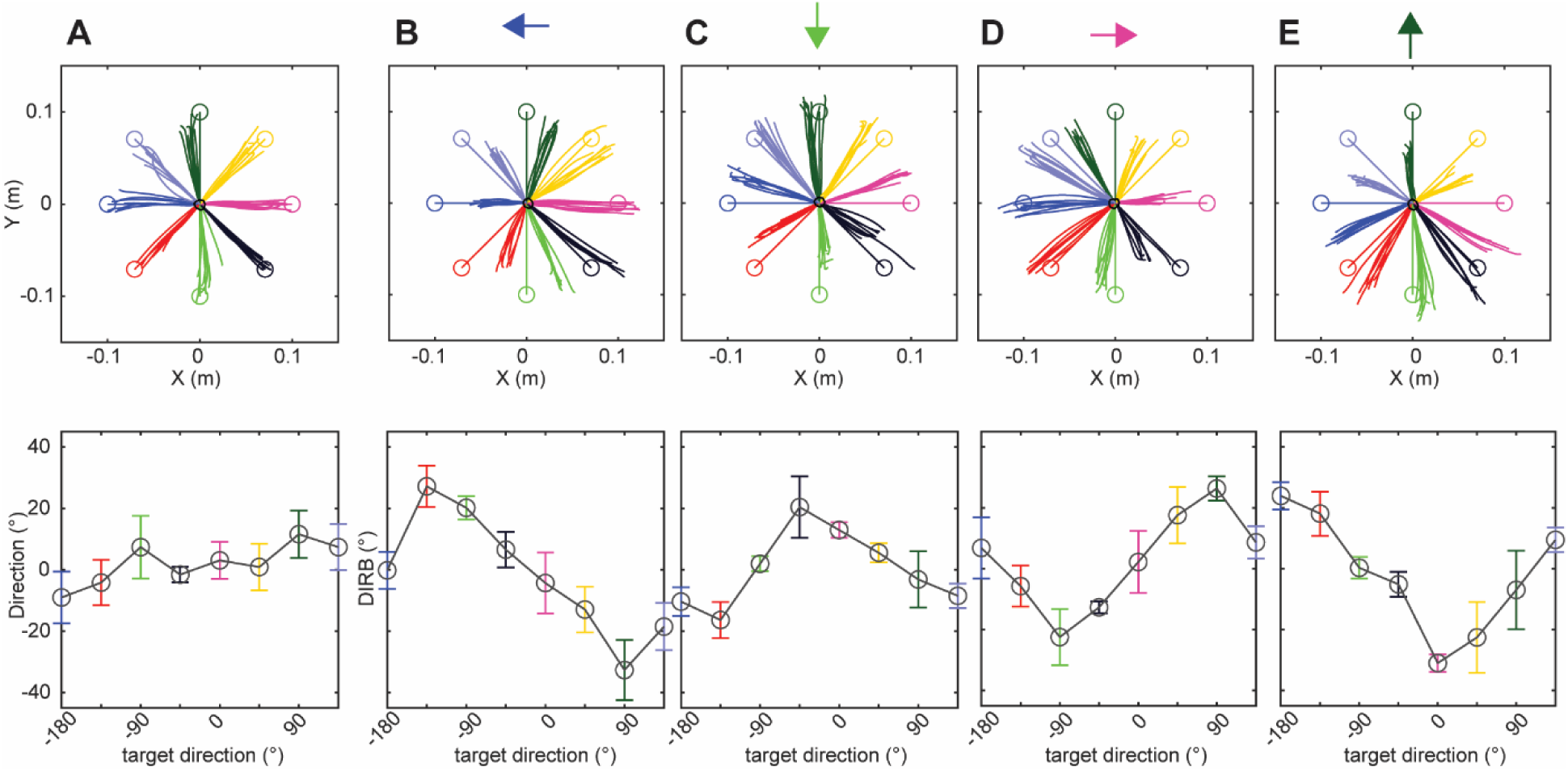
Experimental movement paths (top) and DIRB for a representative subject under different visual shift conditions. A. Movement paths and directions in the no shift condition. B-D: Movement paths and DIRB for medial, posterior, lateral, and anterior visual shifts. Different shift directions resulted in distinct patterns of DIRB as a function of target direction.

**Figure 8.**
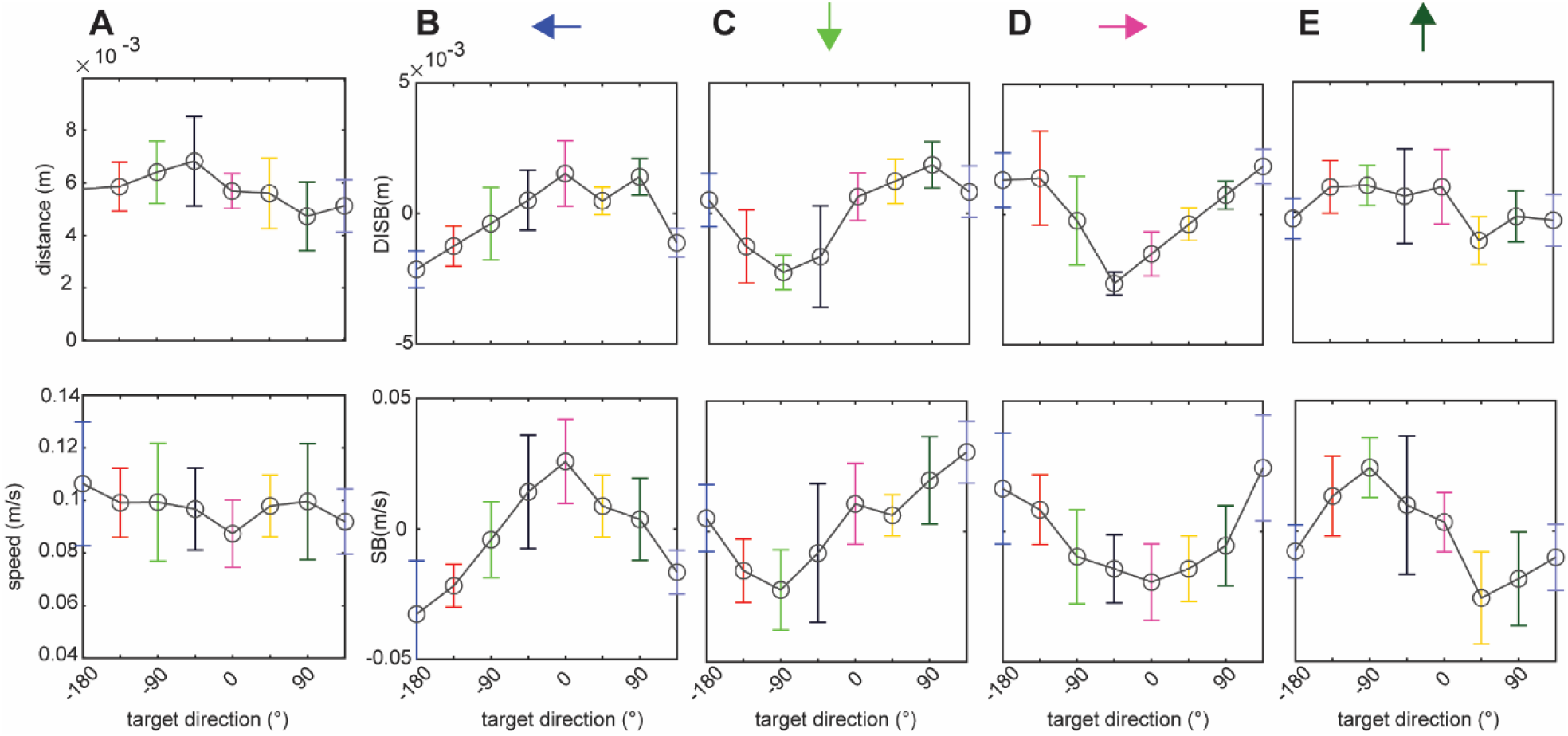
DISB and VB for the subject shown in Fig. 7. A. Movement distances (top) and speeds (bottom) at 40% of peak speed in the no shift condition. B-D: DISB (top) and VB (bottom) for medial, posterior, lateral, and anterior visual shifts. Different shift directions resulted in distinct patterns of DISB and VB as a function of target direction.

Overall, DIRB varied across visual shift directions in a manner that closely followed the predictions of the simulations. Figure 9A shows plots of simulated DIRBs vs. target direction for the four directions of bias used in the experiments. Bias magnitude was fixed in all cases at 0.04 m. For comparison, Fig. 9B shows the subject average (+/-SD) DIRBs in the experiments for the four visual shift conditions. DIRB for the same position bias/visual shift direction showed very similar patterns of variation with target direction. The main differences were regarding the DC offsets of the curves (indicated by the dashed lines), which were obtained by averaging the directional biases across target directions. In the simulated data, these offsets were noticeably different for different position biases, in some cases varying by as much as 10-15°. In the experimental data, differences in average DC bias across visual shift conditions were comparatively small. It should be noted, however, that many individual subjects showed differences in DC bias across shift conditions that were closer in magnitude to those observed in the simulations (data not shown).

**Figure 9.**
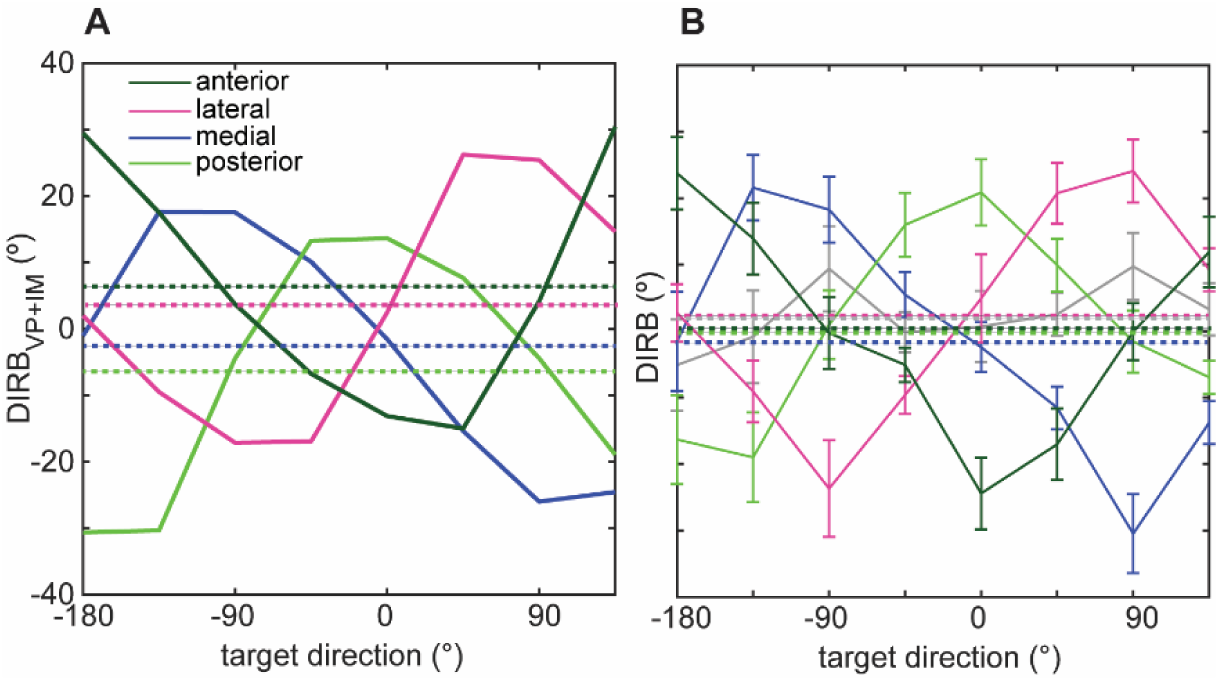
Comparison of simulated and experimental DIRB. A. Simulated DIRB vs. target direction for arm position biases in four directions. Bias magnitudes were 0.04m. B. Subject mean DIRB (+/- SD) vs. target direction for the different experimental visual shift conditions. Gray curve represents the zero-shift condition. Simulated and experimental DIRB varied similarly with target direction for the different directions of bias/shift but curves differed in degree of DC offset.

### 3.6 REAP estimates

REAP estimates derived from directional biases generally conformed to predictions derived from previous studies of somatosensory-visual cue integration employing perceptual paradigms. Figure 10A shows these estimates for all subjects for each of the four visual shifts, superimposed on an image of the hand. In this figure, the center of the hand is assumed to correspond to the brain’s estimated arm position based on somatosensory cues and the ‘X’s indicate the positions of the visual cues. As predicted, REAP estimate distributions for a given visual shift lie in between these positions but are biased more strongly toward the positions of the visual cues, suggesting a primacy of vision in this experimental context (Sober & Sabes, 2003). Moreover, the mean REAP estimates were more biased toward the visual cues for medial and lateral shifts than for anterior and posterior shifts (Table I), in consonance with previous perceptual studies (van Beers et al., 2002). This can also be appreciated from the visual weights in Fig. 10B, which show that the weight distributions for medial and lateral shifts lie closer to a visual weight of one, i.e., the weight if subjects fully relied on vision to estimate their arm positions. These latter trends were confirmed statistically. A one-way repeated measures ANOVA showed a statistically significant effect of shift direction on visual weight (F(3, 57) = [25.097], p<0.001). In addition, post-hoc multiple comparisons revealed that the visual weights did not differ between medial and lateral shifts (Bonferroni adjusted p=0.526) nor did they differ between anterior and posterior shifts (Bonferroni adjusted p=0.613), but did differ between the distributions associated with different axes (Bonferroni adjusted p<0.001).

**Figure 10.**
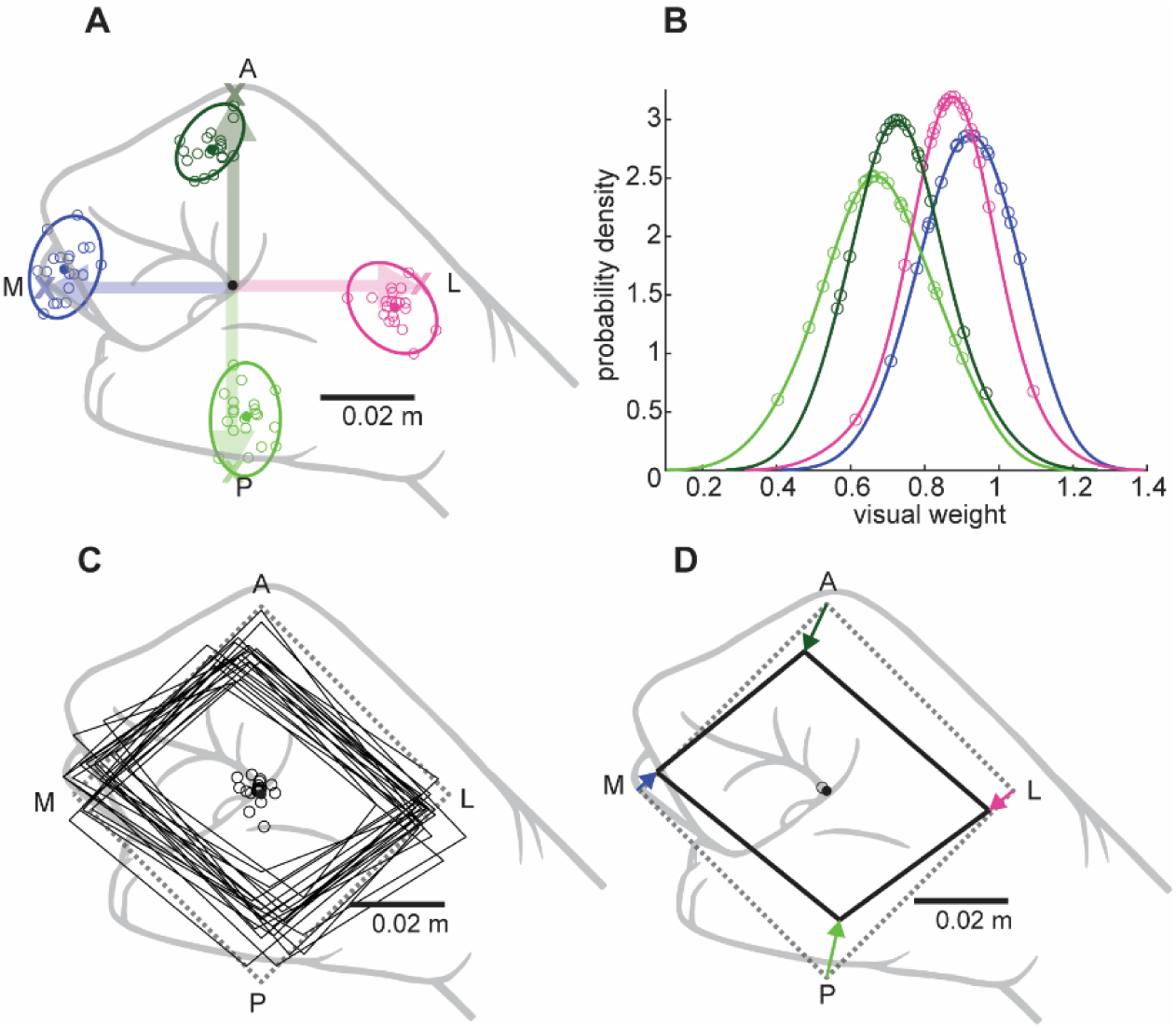
REAP estimates and weights are derived from directional biases. A. Small open circles: individual subject estimates. Filled circles: subject means. Ellipses: 95% confidence intervals for the distributions of subject estimates. ‘X’s: visual cue positions. Arrows: directions and magnitudes of visual cue shifts with respect to presumed somatosensory position cue. B. Probability density functions of the visual weights for the four visual shifts. C-D. Multisensory integration induced distortion of arm position estimates. C. Solid black lines: trapezia formed by connecting each subjects’ REAP estimates across visual shifts. Dotted line: trapezium formed by connecting the visual cues. D. Subject averaged trapezium. Vectors indicate the direction and magnitude of distortion relative to each visual arm position cue.

**Table I.** REAP estimates and associated visual and somatosensory weights.

|  | <u>estimated positions</u> |  |
| --- | --- | --- |
| <u>shift</u> | <u>x (m)</u> | <u>y (m)</u> |
| leftward | -0.0365 +/- 0.0038 | 0.0041 +/- 0.0054 |
| backward | 0.0028 +/- 0.0037 | -0.0277 +/- 0.0053 |
| rightward | 0.0348 +/- 0.0037 | -0.0042 +/- 0.0039 |
| forward | -0.0048 +/- 0.0035 | 0.0299 +/- 0.0039 |
| no shift | 0.0036 +/- 0.0048 | -0.0009 +/- 0.0051 |
|  | <u>weights</u> |  |
| <u>shift</u> | <u>visual</u> | <u>somatosensory</u> |
| leftward | 0.911 +/- 0.0944 | 0.088 +/- 0.0945 |
| backward | 0.691 +/- 0.1320 | 0.308 +/- 0.132 |
| rightward | 0.8698 +/- 0.0933 | 0.1302 +/- 0.0933 |
| forward | 0.7468 +/- 0.0984 | 0.2532 +/- 0.0983 |

Multisensory integration led to an anisotropic distortion of subjects’ arm position estimates. Figure 10C shows a visual representation of this distortion for each subject, obtained by joining each subject’s REAP estimates across visual shifts. Compared to the trapezium formed by joining the positions of the visual cues (dotted lines), the individual REAP estimate trapezia appear contracted toward the center of the hand, but to a larger extent for cues on the anteroposterior or ‘depth’ axis than for those on the horizontal axis, consistent with the analyses of the visual weights. In addition, for most subjects the vertices of the trapezia were also shifted away from the vectors joining the somatosensory and visual cues, giving the trapezia a ‘twisted’ appearance. Stated differently, the position estimates had both axial and off-axis components, both of which depended on the direction of visual shift. This can be appreciated from Fig. 10D, which shows the magnitude and direction of distortion for each average position estimate. Average off-axis components were significantly different from zero for all shifts, though these differences were somewhat larger for the anterior (t = −6.175, df = 19, p<0.001) and lateral (t = −4.840, df = 19, p<0.001) shifts, than for the posterior (t = 3.367, df = 19, p=0.003) and medial (t = 3.374, df = 19, p=0.003) shifts.

A follow-up experiment (Experiment 2) provided strong support for the assertion that REAP estimates reflect the hand positions used during motor planning. Figures 11A-B illustrate the rationale behind this experiment. Figure 11A shows simulation results obtained under conditions where arm position was biased 0.04 m lateral and targets were in their standard positions. Under these conditions, arm position bias led to substantial DIRB (see also Fig. 9). Figure 11B shows simulated DIRB for the same arm position bias but under conditions where the targets were shifted to match the arm bias direction and magnitude, i.e., they were centered on the biased position of the hand. Under these conditions DIRB are predicted to be greatly reduced, due to the elimination of VP errors. Some DIRB are still expected, however, due to IM errors, which persist because of the lingering difference between the actual and biased arm positions. Overall, these simulations predict that if target positions are shifted such that they are centered on subjects’ REAP estimates rather than their actual hand positions, then DIRB should be strongly attenuated.

**Figure 11.**
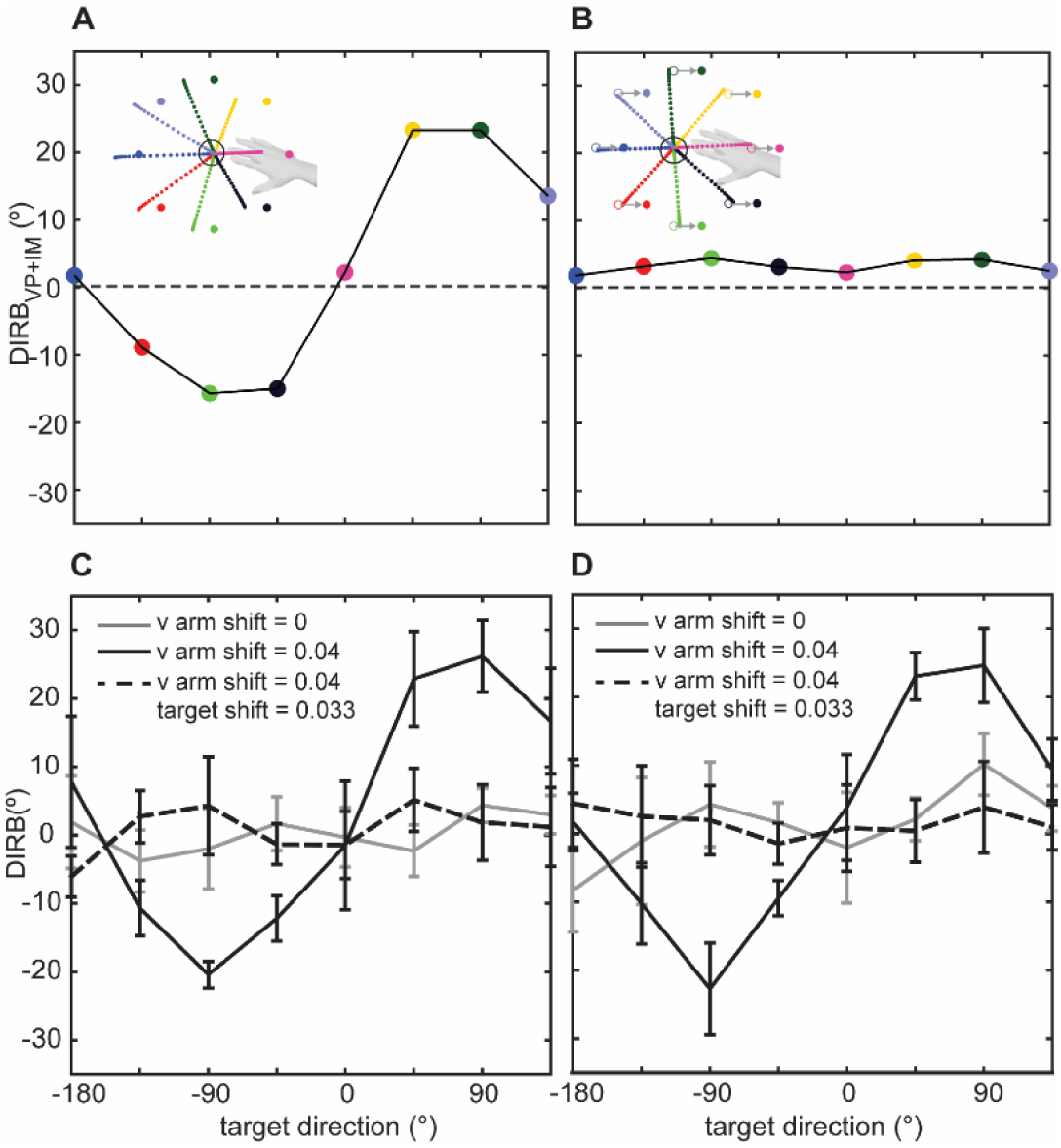
Directional biases under unshifted and shifted target conditions. A. Simulated DIRB when arm position is biased (0.04m lateral (0°)) and targets are unshifted. B. Simulated DIRB for the same arm position bias but when targets are shifted to match the bias direction and magnitude. DIRB are predicted to be largely attenuated. C. Experimental DIRB for a single subject under unshifted and shifted target conditions. Grey curve: mean trial DIRB (+/- SD) under conditions of no arm cue shift and no target shift. Solid black curve: DIRB for a 0.04m lateral arm cue shift and no target shift. Dashed black curve: DIRB for a lateral arm cue shift (0.04 m) combined with a lateral target shift (0.033m). When targets were shifted to match the expected REAP estimate, DIRB were largely attenuated. D. Same format as A but showing means and SDs over subjects (N=6). In all panels, target direction is defined with respect to the unshifted target directions.

Figures 11C-D show that DIRB in the presence of combined arm position bias and target shifts were greatly reduced, consistent with the simulation predictions. Figure 11C shows the results of the follow-up experiment for a single subject. The grey curve shows DIRB in the absence of imposed arm bias and target shifts. Here, DIRB were negligible, as expected. The solid black curve shows DIRB when the visual arm position cue was shifted lateral by 0.04m, which led to substantial DIRB for some target directions (see also Figs. 9 and 7). The dashed black line shows DIRB when, unbeknownst to the subjects, in addition to this imposed arm position bias, target positions were shifted such that they were centered on the average REAP estimate for lateral visual shifts in Experiment 1 (i.e., 0.03m). Under these conditions DIRB were greatly reduced and were similar in magnitude to those observed in the no-shift condition. Figure 11D shows the average DIRB for six subjects tested under the same three conditions, which show the same basic trends. The results of this experiment provide strong support for the idea that hand position estimates derived from the REAP method reflect those that were used during motor planning in these experiments.

REAP estimates derived from speed biases were highly variable and less in line with predictions based on previous perceptual studies than those derived from DIRB. Figure 12A shows the SB-derived estimates for all subjects for each of the four visual shifts, in the same format as Figure 10A. Although the means of these position estimate distributions were biased in the directions of the visual cues (as expected) the magnitudes of these biases were generally less than those derived from DIRB and were more uniform across shift directions. Moreover, position estimates were much more variable across subjects than those derived from DIRB, which likely reflects the fact that peak movement speeds were also highly variable across subjects. In addition to being more variable, estimates for some directions, e.g., medial and posterior, were somewhat irregularly distributed in space, which was reflected in the distribution of visual weights (Fig. 12B). The large variability in REAP estimates across subjects can also be appreciated from Fig. 12C, which shows the trapezia obtained by joining the position estimates of each subject. This figure, as well as the subject average trapezium (Fig. 12D) suggests that REAP estimates derived from SB were less influenced by the visual cues than those derived from DIRB. That is, these trapezia showed more contraction toward the presumed position of the somatosensory cues and noticeably less twisting of the vertices than those shown in Fig. 10. The potential significance of this finding is discussed below.

**Figure 12.**
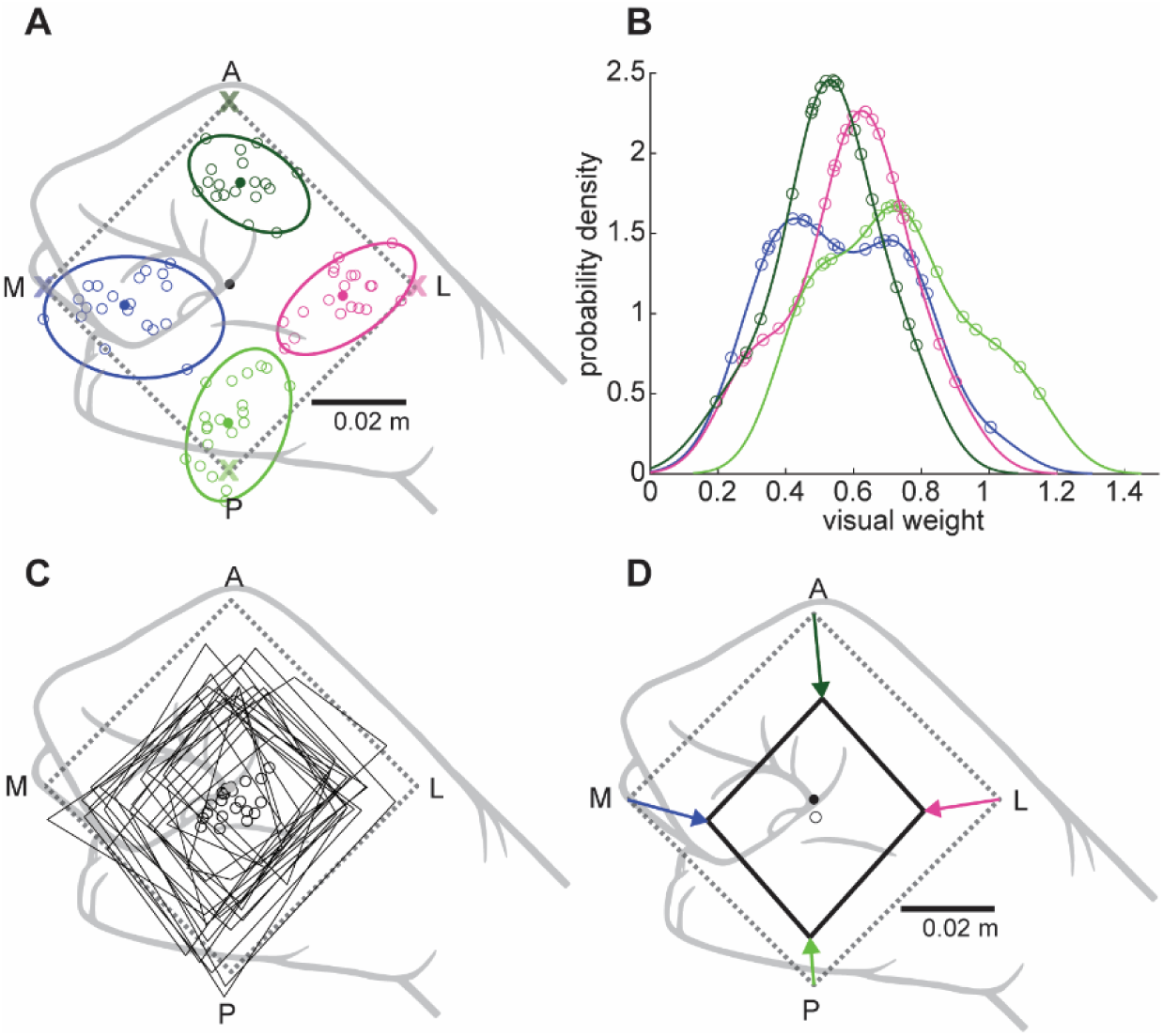
REAP estimates and weights derived from speed biases. Format as described in Fig. 10.

## 4 Discussion

In this study, we combined human psychophysical experiments, forward dynamic simulations, and mathematical optimization to reverse engineer arm positions (‘REAP’) from multisensory integration induced motor biases. We hypothesized that these REAP estimates would mirror characteristics of multisensory position estimates and weights reported in previous studies employing perceptual tasks. Position estimates derived from biases in initial movement directions (DIRB) largely supported this hypothesis in that they reflected the integration of both visual and somatosensory cues but were more influenced by vision for cue shifts along the horizontal axis than for shifts along the anteroposterior axis. However, these estimates did not fall on the straight lines connecting the visual and somatosensory arm position cues, suggesting that a simple linear weighting scheme may be inadequate for describing cue integration during motor planning. Estimates derived from speed biases were more variable and less in line with hypotheses based on perceptual experiments. Regarding the DIRB-derived estimates, the similarities between the present results and those obtained in perceptual experiments have important implications for understanding arm proprioception, as they suggest that the brain’s perceptual and action systems use largely similar mechanisms to estimate arm position from multisensory cues. The present results could also have important implications for clinical assessments of position sense. That is, they suggest that assessments derived from motor biases can be used to augment perceptual assessments or be used in place of them when perceptual reporting is impractical or impossible.

### 4.1 REAP estimates from directional biases: positions and visual weights

Several previous studies have reported directional and workspace variations in arm position sensing. For example, van Beers and colleagues studied the ability of human subjects to localize visual or somatosensory proprioceptive targets at three positions in the horizontal plane (van Beers et al., 1998). They found that subjects were more precise (i.e., less variable) when localizing positions along an anteroposterior axis than along a mediolateral one. In addition, subjects were more precise when localizing positions closer to the body than farther away. Subsequent work employing visuomotor adaptation paradigms (van Beers et al., 1999; van Beers et al., 2002) confirmed the direction-dependent precision of both somatosensory and visual localization and described some of the basic rules underlying the integration of these cues for position estimation. The findings for somatosensory-based localization were later confirmed in studies employing robots to assess proprioceptive abilities in a horizontal plane (Wilson et al., 2010) and, more recently, in 3D space (Klein et al., 2018). However, although these studies provided important information regarding the accuracy and precision of proprioceptive estimates, they did not specifically address proprioception’s role in position estimation during motor planning.

Studies by Sober and Sabes directly addressed how somatosensory and visual feedback were combined to estimate arm position during motor planning (Sober & Sabes, 2003, 2005). They showed that these two sensory signals were weighted differently during the VP and IM stages and that these weights also varied with the sensory modality used to cue target position and with the information content of the visual cue (fingertip vs full arm vision). Since the focus of their studies was on multisensory integration and not position sensing per se, position estimates were not explicitly reported. However, based on the reported weights, it can be inferred that these estimates were strongly biased toward the positions of the visual cues in their experiments. Based on these results and the aforementioned perceptual studies, we hypothesized that position estimates derived from motor biases (which were assumed to reflect the integration of cues during motor planning) would generally be biased more towards the positions of the visual cues. Additionally, based on other work (van Beers et al., 1998, 1999; van Beers et al., 2002) we hypothesized that the extent of this bias would depend upon the direction of the visual shift, i.e., estimates would be more biased towards the positions of the visual cues for medial and lateral visual shifts than for anterior and posterior shifts.

Our results confirmed these hypotheses. Position estimates were generally biased toward the positions of the visual cues. In addition, the extent of this visual bias was axis dependent; estimates were more strongly biased toward the visual cues for shifts along the horizontal axis than along the anteroposterior (or depth) axis. However, similar to Sober and Sabes, we assumed that position estimates would evolve from a straightforward weighted sum of somatosensory and visual cues and would thereby lie along the straight line connecting the positions of these cues. This was not observed: average position estimates had components that were both on-axis, i.e., collinear with the visual shift axes, and off-axis components, giving rise to the twisted appearance of the trapezia shown in Fig. 10. The source of these off-axis components is currently unclear. One possibility is that they arise from methodological factors. Among these was our decision to calculate experimental directional biases with respect to the initial movement directions in the no-shift condition, rather than with respect to the cued target directions. When directional biases were calculated with respect to these latter directions, average position estimates were similar but more variable across subjects (Fig. S4). In addition, many individual subjects still demonstrated position estimates with substantial off-axis components, though on average these were reduced somewhat for some shift directions (e.g., medial and lateral) or differed in direction (e.g., for anterior shifts) relative to those calculated with respect to the no-shift condition. As a result, the extent to which this methodological factor contributed to the occurrence of off-axis position components is equivocal.

Other factors related to the nature of visual and somatosensory position cues and their integration could have contributed to the presence of the off-axis position components. For example, van Beers and colleagues have shown that a computational model employing visual and somatosensory arm position cues that are weighted according to their direction-dependent precision predicts that integrated position estimates should (in at least some instances) lie off the straight line connecting the two cues, a prediction that was confirmed experimentally (van Beers et al., 1999). It is conceivable, therefore, that the ‘off-axis’ components of the position estimates observed in the present study evolved at least in part from a precision-dependent weighting of visual and somatosensory position cues. Another more recent model (Wang, 2024) posits that motor biases results from systematic errors in transforming hand and target position from visual coordinates (Buneo & Andersen, 2006; Buneo et al., 2002) to proprioceptive coordinates (Helms-Tillery et al., 1991). Stated somewhat differently, motor biases are believed to be caused primarily by a ‘misalignment’ between eye-centered and body-centered representations of position. It is not difficult to imagine how such a misalignment could result in integrated position estimates that lie off the axes of the visual shifts in the current experiments, though additional experiments and more in-depth analyses are required to reconcile the predictions of this model with the present results.

### 4.2 Directional biases: DC offsets

In this study, when simulated DIRB were plotted as a function of target direction, the resulting curves differed in their DC offsets for different visual shifts, a finding that was not observed in the experimental data. In the simulations, DC offsets differed by as much as 12° or more between visual shifts, while DC offsets were negligible in the population data from the experiments. The basis for these differences is currently unclear, but it is important to reiterate that the lack of prominent DC offsets in the population data did not substantially influence the main results. The REAP procedure was performed using subjects’ individual motor biases, and, as mentioned in the Results, many individual subjects showed substantial differences in DC offsets across shift directions. Moreover, removing the DC offsets from the simulation data by subtracting the mean DIRB (across target directions) affected the position estimates and weights only minimally. Another possibility is that the differences in DC offsets relate to differences in arm posture between the simulations and the experiments. In the simulations, the arm was constrained to lie in the horizontal plane of the targets. However, due to space limitations imposed by the apparatus, in the experiments subjects’ arms were rotated into a posture that was more vertical. As shown in Fig. 5, changing the arm posture within the plane of the targets did not change the target dependence of DIRB but did change the DC bias. Thus, it is entirely possible that the relatively small DC biases observed in the experiments could simply be due to the arm posture that was used. Importantly, there is one additional and important difference between the simulations and experimental data that could account for the differences in DC offsets.

As noted in the Methods, the simulations were entirely feedforward, i.e., no somatosensory or visual feedback was included. For the experiment, this was obviously not the case, so it is entirely possible that the lack of DC offsets can be explained by the presence of feedback, particularly somatosensory feedback. As noted in the Results, the DC component of the DIRB can be traced to errors occurring at the IM stage of motor preparation, where somatosensory input is thought to be weighed more strongly than visual input (Sober & Sabes, 2003). Thus, the lack of a significant DC bias in the experimental data could reflect a role for somatosensory feedback in partially attenuating position estimation errors that would otherwise result from mismatches between visual and somatosensory cues.

### 4.3 REAP estimates from speed biases

Although our focus was on deriving arm position estimates from biases in movement direction, we also investigated using speed biases to derive position estimates. Note that we could have also used distance biases, as distance and speed were highly correlated in the experiments (Fig. S3). Nevertheless, results using speed biases were similar in some ways to those using directional biases in that average position estimates were clearly shifted in the directions of the visual cues. However, estimates derived from speed biases differed in several ways from those derived from direction. First, they were much more variable across subjects (Fig. 12A). This could be a consequence of the fact that peak movement velocities were, in general, quite variable across subjects. However, even after normalizing each subject’s velocity data, estimates were still considerably more variable than those derived from directional biases. This variability was also reflected in the distributions of visual weights, which were not normally distributed for all shift directions (Fig. 12B). The larger variability in the position estimates can also be appreciated from subjects’ individual trapezia (Fig. 12C), though there were still some general trends that can be gleaned from these and the average trapezium. First, position estimates did not have significant off-axis components and therefore the trapezia appeared less twisted than those derived from directional biases. In addition, average position estimates were less biased by the visual cues and, as a result, the average trapezium showed a larger degree of contraction toward the presumed position of the somatosensory cues than the one derived from directional biases. Given the relatively large variability in the estimates, it is difficult to draw definitive conclusions about these summary findings. However, if the findings are confirmed in future experiments, it could indicate that position estimates for the specification of movement distance/speed differ from those used for the specification of movement direction, and that the former are more reliant on somatosensory cues than those for direction. This would provide additional support for the idea that movement direction and extent are planned independently (Gordon, Ghilardi, & Ghez, 1994; Sainburg et al., 2003; Vindras et al., 1998). Future experiments employing more stringent speed demands could help clarify these issues.

### 4.4 Clinical Implications

Proprioceptive dysfunction is a known sequela of several neurological disorders. In most clinical environments, proprioceptive assessments are still performed manually by clinicians. Although these assessments can be performed quickly, they suffer from several drawbacks. First, the qualitative nature of these assessments is associated with poor or modest inter- and intra-rater reliability (Carey, 1995; Lincoln et al., 1991) and can be difficult to track over time. In addition, since these tests involve manually guiding a subject through a movement or placing the limb in position, assessments can be contaminated by physical cues transmitted by the clinician (Simo et al., 2014). The introduction of robotic assessment systems has alleviated many of these issues, but current assessment protocols are still largely incomplete in that they only directly address proprioception’s perceptual components. That is, one of the two major neuroanatomical pathways carrying proprioceptive information arises from the muscles and tendons and ends in the somatosensory cortex. This pathway is believed to underlie our ability to recognize and verbally report the position of our limbs in space, i.e., ‘conscious’ proprioception. However, proprioception derived from the muscles and tendons is also processed in pathways that terminate in the cerebellum. Information traveling in this pathway is processed at a largely unconscious level but is believed to be particularly important for motor control. It is unclear, therefore, the extent to which methods used to assess *conscious* proprioception can capture the information carried by this pathway. The REAP protocol described here can, in principle, help to fill this important gap in proprioceptive assessments, though several refinements will be required before it can be incorporated into existing clinical protocols. These refinements include minimizing the number of trials and/or movement directions used to derive motor biases. In addition, given that some of the between-subject variability in estimates, sensory weights, and DC offsets observed here could have arisen from the use of generic anthropomorphic data for the modeling and simulation methods, future refinements could include customizing these procedures by incorporating each subject’s own anthropomorphic data. In addition, to be maximally useful, the protocol should be tested and validated under conditions without visual shifts, to allow for assessment of position sense when proprioceptive information is provided solely by somatosensory cues.

The REAP method described here also has the potential to impact the development of upper extremity neural prostheses. A current major focus of research in this field is aimed at endowing these devices with sensory feedback, including proprioception, with the goal of fostering embodiment. Successful embodiment of devices is thought to be dependent upon both an awareness of ownership (the sense that the device is part of our own body) and agency (the sense of control over the actions of the device). An improved understanding of position sensing in the context of motor planning may provide important insights into how artificial proprioceptive signals could be optimally structured to support embodiment, particularly agency. More specifically, our methods for quantifying the relation between position estimation and movement errors could be useful for assessing whether artificial proprioceptive signals are being appropriately integrated during arm position sensing and movement planning. In addition, an improved understanding of unconscious proprioceptive processing could potentially lead to the development of artificial feedback systems that mitigate the effects of cognitive burden on the user, a known yet unresolved issue with traditional artificial feedback.

## 6 Author Contributions

KA: Data curation, Formal analysis, Investigation, Methodology, Software, Validation. HM: Data curation, Formal analysis, Investigation, Methodology, Software, Validation. HL: Methodology, Validation, Writing – review & editing. CB: Conceptualization, Data curation, Formal analysis, Investigation, Methodology, Software, Validation, Visualization, Writing – original draft, Writing – review & editing, Funding acquisition, Project administration, Resources, Supervision.

## 8 Acknowledgments

The authors would like to thank Dr. Marco Santello for the use of the KINARM system and for his feedback on various aspects of the project. The authors would also like to thank Dr. Stephen Helms Tillery for his valuable feedback on an initial version of the manuscript.

