## Supplementary figures and images for "Arm position estimates derived from motor biases"

### Supplemental Figure 1

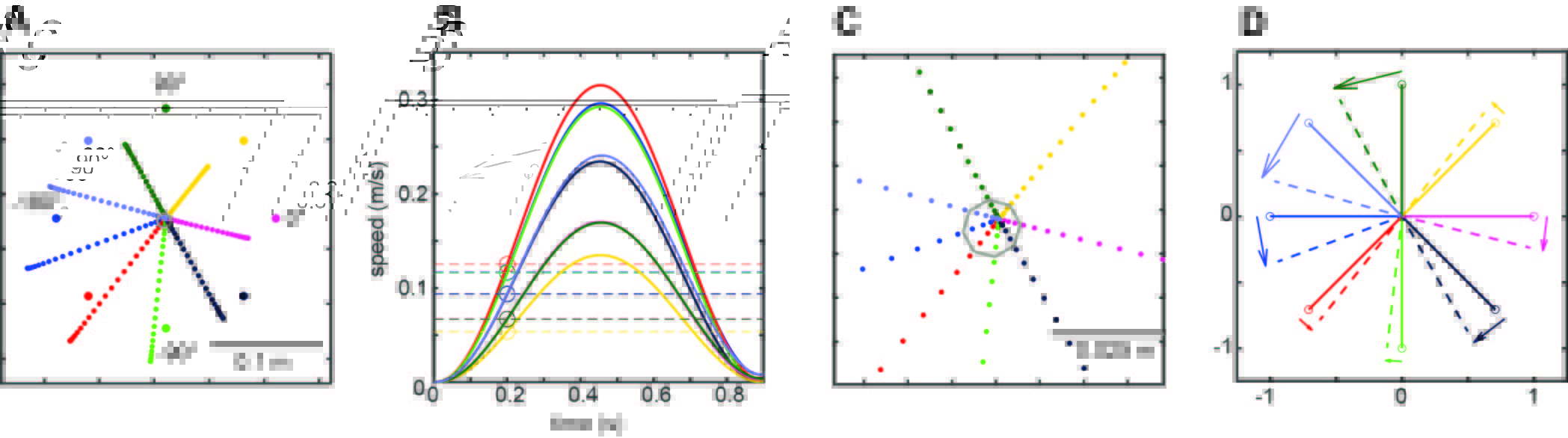

### Supplemental Figure 2

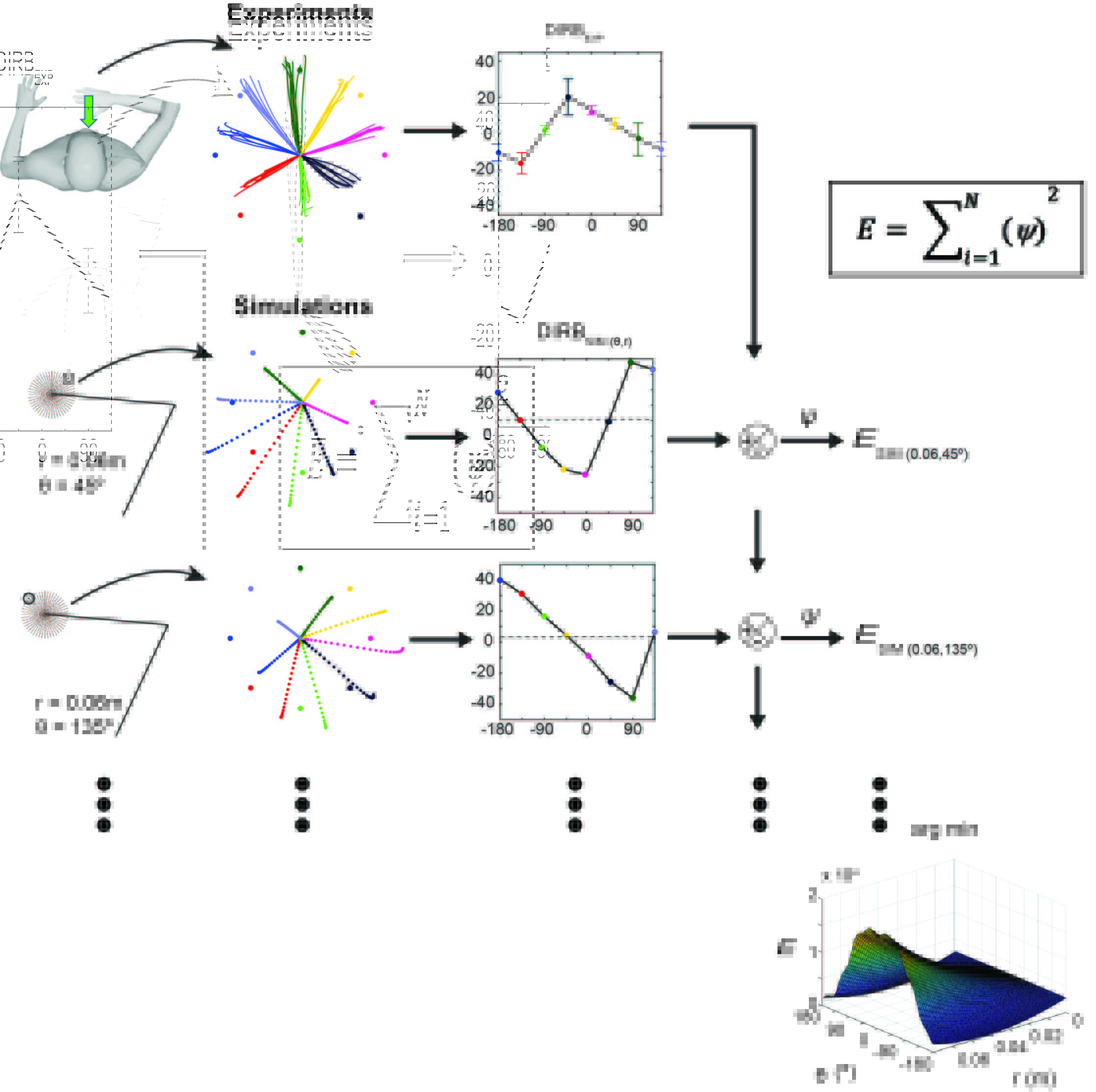

### Supplemental Figure 3

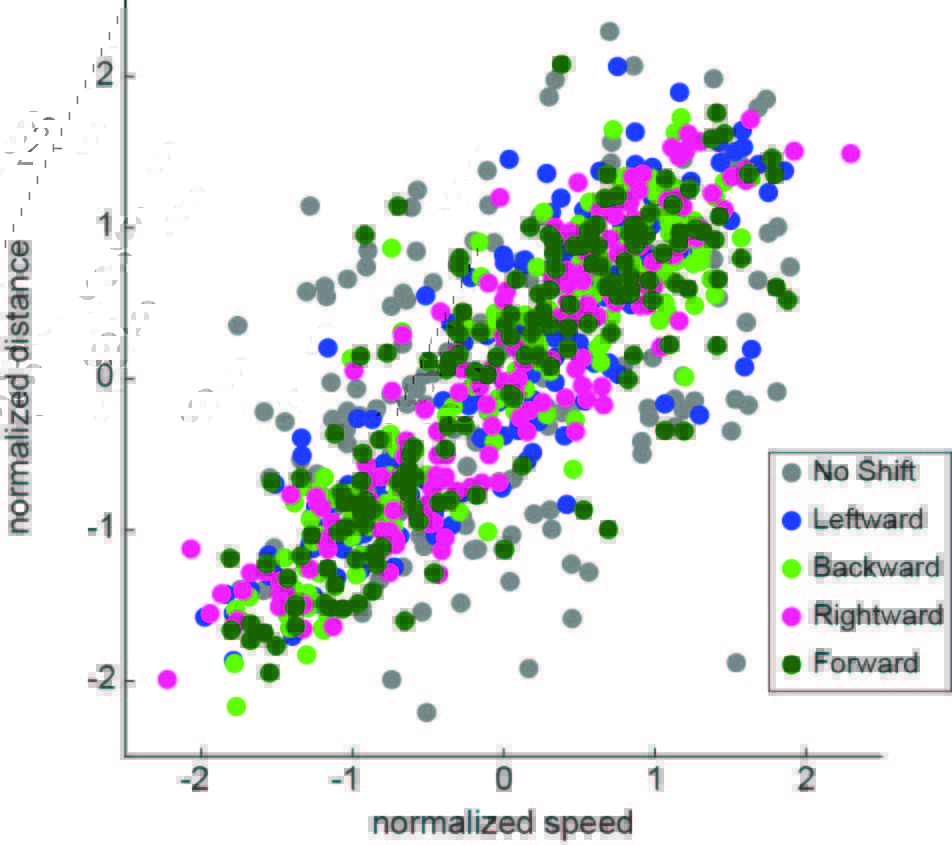

### Supplemental Figure 4

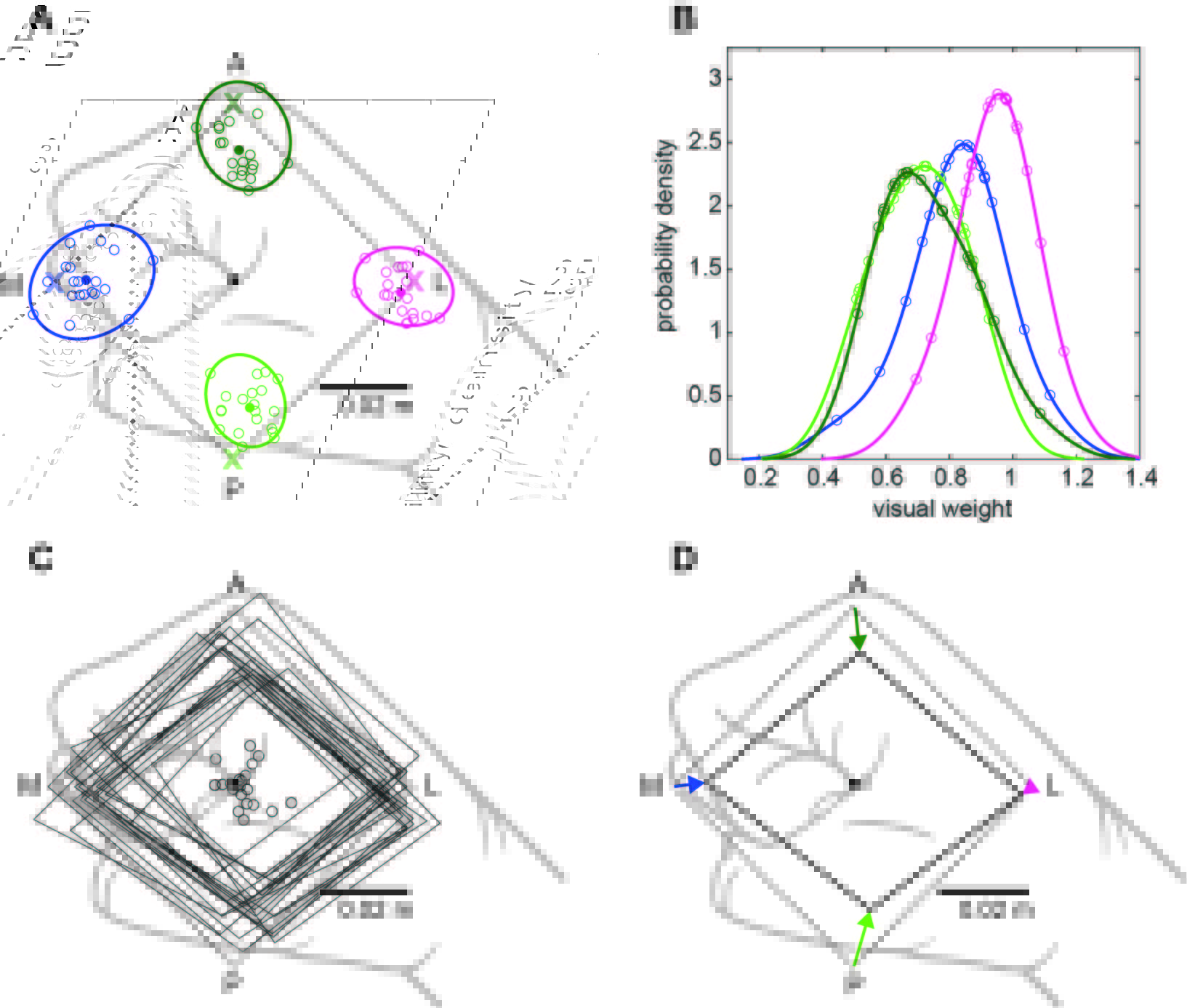
